# Nav1.8 mediates peripheral-to-central nociceptive transmission independently of central presynaptic mechanisms in human DRG-spinal cord circuits

**DOI:** 10.64898/2026.02.27.708640

**Authors:** Seph M. Palomino, Katherin A. Gabriel, Ashwin Koduri, Ibrahim Khan, Rahman Syed A, Peter Horton, Tariq Khan, Anna Cervantes, Geoffrey Funk, Stephanie Shiers, Theodore J. Price, Amol Patwardhan

**Author notes:** Authors contributed equally to this work (Seph M. Palomino, Katherin A. Gabriel). **Declaration of Interests:** Conflict of Interest Statement: T.J.P. is a co-founder of and holds equity in 4E Therapeutics, NuvoNuro, PARMedics, and Nerveli. T.J.P. has received research grants from AbbVie, Eli Lilly, Grunenthal, Evommune, Hoba Therapeutics, and The National Institutes of Health. The authors declare no conflicts of interest related to this work.

## Abstract

Suzetrigine, a selective Nav1.8 blocker, despite its significant role as a non-opioid analgesic, exhibits modest efficacy. The potential enhancement of Nav1.8 blocker efficacy through engagement of Nav1.8 at central terminals remains uncertain. We established a donor-derived human organotypic nociceptive circuit to determine where and how clinically relevant analgesics suppress pain signaling. Using ELISA and immunohistochemistry, we quantified calcitonin gene-related peptide (CGRP) release from acute explants of adult human spinal cord, dorsal root ganglia (DRG), and an intact DRG-spinal cord preparation preserving primary afferent anatomy and directional signaling. In isolated spinal cords, capsaicin evoked concentration-dependent spinal CGRP release without compromising viability, and clinically used analgesics inhibited it, validating the assay. In the intact circuit, the application of capsaicin to the DRG cell bodies triggered CGRP release exclusively in the spinal cord, consistent with compartmentalized neuropeptide release at central terminals. Selective Nav1.8 inhibition with suzetrigine reduced spinal CGRP release only when applied to the DRG or nerve root, not the spinal cord, indicating that Nav1.8 regulates peripheral action potential propagation rather than presynaptic transmitter release. These findings establish the first intact human pain circuit-based assay to study analgesics and demonstrate that the analgesic efficacy of Nav1.8 inhibitors is unlikely to increase with improved CNS penetration.

## Introduction

Chronic pain is a major public health burden, affecting roughly one in five American adults, with about 10% of them suffering from high impact chronic pain that limits daily function (1). Despite extensive preclinical target discovery efforts, translation of analgesic mechanisms from animal models to effective human therapies has remained challenging. Few experimental systems permit direct testing of pharmacologic modulation within intact human nociceptive circuits, and developing such platforms is critical for mechanistic validation of emerging non-opioid analgesics.

Nociceptive neurotransmission depends on the propagation of nociceptor action potentials from the periphery to the spinal dorsal horn, where neuropeptide release regulates key aspects of synaptic transmission (2). The neuropeptide calcitonin gene-related peptide (CGRP) is abundantly expressed in TRPV1-positive nociceptive dorsal root ganglion (DRG) neurons and is released from their central terminals in the spinal dorsal horn. CGRP contributes to both peripheral neurogenic inflammation and central sensitization, and its signaling has been implicated across multiple chronic pain states, including migraine and inflammatory pain (3, 4).

Nociceptor excitability is controlled by specialized voltage-gated sodium channels, including Nav1.8, which is selectively expressed in small-diameter sensory neurons (5–7). In animal models, genetic, antisense oligonucleotide, and pharmacological inhibition of Nav1.8 reduces nociceptor hyperexcitability in sensory neurons and alleviates pain behaviors in both inflammatory and neuropathic pain models (8–11). This expression profile and in vivo efficacy data in animals established Nav1.8 as a validated analgesic target, and motivated the discovery and clinical development of JOURNAVX (suzetrigine), a selective Nav1.8 inhibitor (12–18), that was recently approved by the FDA for the management of moderate to severe acute pain (19). Suzetrigine has demonstrated clinical efficacy in acute pain; however, its efficacy across chronic pain conditions has yet to be established.

Pan sodium channel inhibitors such as lidocaine and bupivacaine produce robust spinal analgesia when delivered intrathecally but cause dose-limiting motor and central nervous system side effects due to nonselective channel blockade (20, 21). Because Nav1.8 is largely restricted to peripheral nociceptors, selective inhibition of this channel could, in principle, provide sensory-selective analgesia when applied at spinal levels. However, it remains unknown whether Nav1.8 localizes along centrally projecting human DRG axons, contributes to action potential propagation in intact human circuits, or regulates neurotransmitter release from central terminals. Moreover, while Nav1.8 is important for human nociceptor excitability, other voltage-gated sodium channels such as Nav1.7 are also expressed in human DRG neurons and their relative contribution in a human pain circuit is unknown (14–16). This raises the question of whether selective Nav1.8 blockade alone is sufficient to suppress central nociceptive output.

In this study, we establish complementary human organotypic preparations of spinal cord, DRG, and an intact DRG-spinal cord circuit that preserves directional peripheral-to-central connectivity. Using TRPV1 stimulated-CGRP release as a quantitative functional readout of nociceptive signaling, we assess how pharmacologic modulation at distinct anatomical sites governs central neuropeptide output. By combining these human peripheral-central circuit preparations with selective Nav1.8 inhibition, we define the anatomical domain along the human nociceptive axis where Nav1.8 is likely controlling signal propagation leading to central neuropeptide release. These studies provide mechanistic insight into the spatial basis of Nav1.8 dependent analgesia and establish a human-specific platform for evaluating analgesic mechanisms within intact nociceptive circuitry.

## Results

### Human DRG-spinal cord circuit restricts CGRP release to central terminals

To interrogate human nociceptive signaling, we developed an acute organotypic DRG, spinal cord, and DRG-spinal cord explants preparations from organ donor-derived tissues, preserving primary afferent connectivity and directional peripheral-to-central CGRP signaling (Figure 1A-F). In human DRG, TRPV1 is broadly expressed across many nociceptors, and a large subset of these neurons co-express CGRP (22–24); accordingly, capsaicin was used to selectively activate TRPV1-positive afferents and evoke neuropeptide release. Isolated spinal cord slices alone exhibited concentration-dependent CGRP release in response to capsaicin, with 10 µM inducing a 2.5-fold increase (250%), 5 µM a 1.7-fold increase (170%), and 1 µM a 1.2-fold increase (119%) relative to baseline (100%) (Figure 1B, D). Similarly, isolated DRG slices alone exhibited CGRP release in response to 1 µM capsaicin at a 1.75-fold increase (175%) relative to baseline (Figure 1D). Viability assays confirmed that capsaicin-induced CGRP release was not due to tissue toxicity (Figure 1C, E).

**Figure 1.**
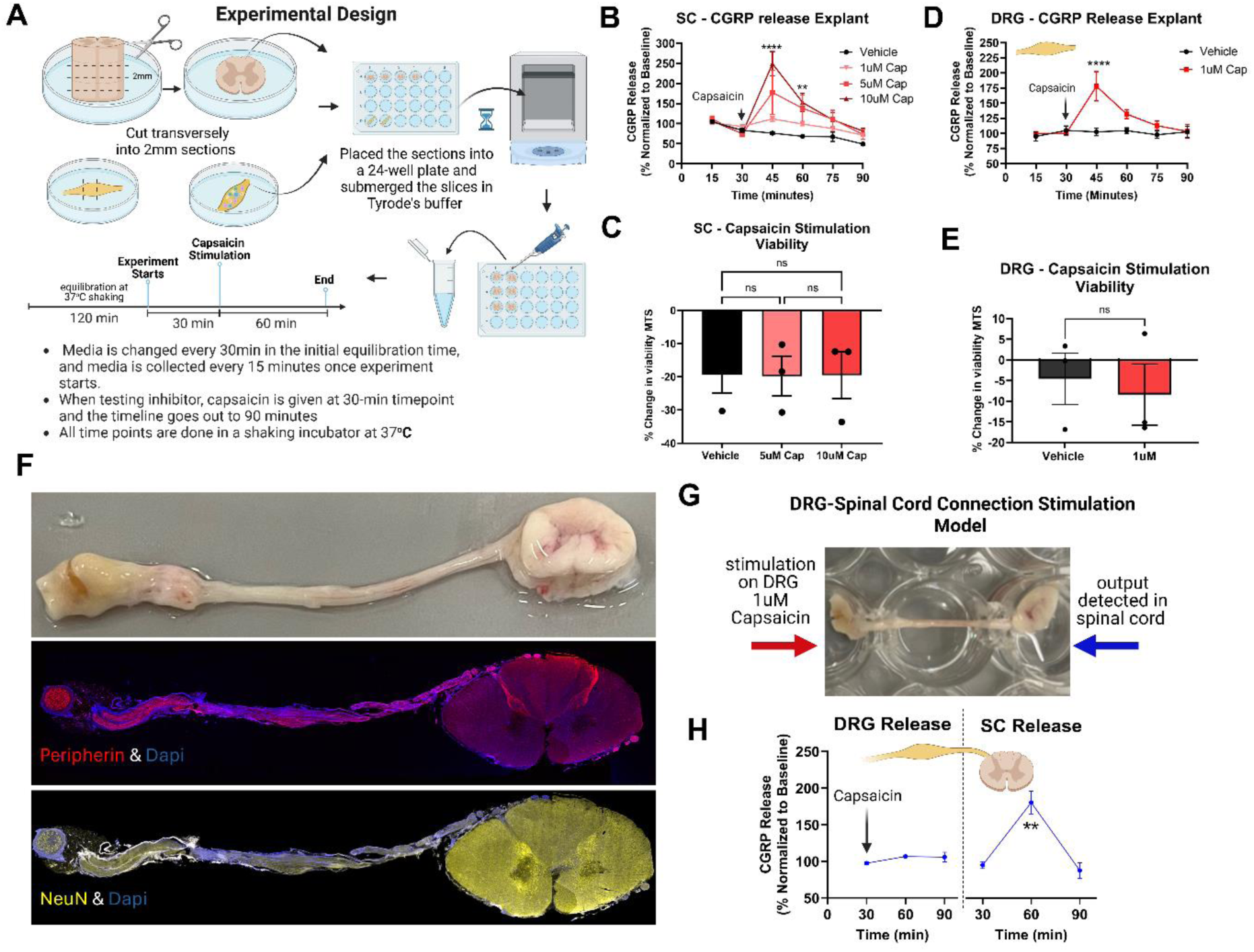
Organotypic human DRG-spinal cord circuit restricts CGRP release to spinal central terminals. An intact human DRG-spinal cord explant reveals that nociceptor activation evokes CGRP release selectively at spinal central terminals, not from the DRG soma. Acute adult human spinal cord explants were used to establish a quantitative, activity-dependent assay of central CGRP release in response to nociceptor activation. **(A**) Experimental workflow describing the ex-vivo spinal cord and dorsal root ganglion preparation, conditions and timeline for the capsaicin evoked CGRP release. (**B**) Time course of CGRP release in response to increasing concentrations of capsaicin (1µM, 5µM, 10µM) showing a concentration-dependent increase in the magnitude of release. Data are means ± SEM; n = 3-4 individual donors. **P < 0.01, and **** P < 0.0001 versus same time point in the Vehicle group by repeated measures (RM) two-way ANOVA with Tukey’s post hoc test. (**C)** MTS % change in viability in spinal cord explants treated with capsaicin (5µM or 10µM) for 30 minutes. There were no statistically significant differences between any of the groups using a one-way ANOVA and Tukey’s multiple comparison test. (**D)** Time course of CGRP release in response to capsaicin (1µM) showing increase in the magnitude of release. Data are means ± SEM; n = 3-4 individual donors. **** P < 0.0001 versus same time point in the Vehicle group by repeated measures (RM) two-way ANOVA with Tukey’s post hoc test. **(E)** MTS % change in viability in spinal cord explants treated with capsaicin (5µM or 10µM) for 30 minutes. There were no statistically significant differences between any of the groups using a one-way ANOVA and Tukey’s multiple comparison test. **(F**) Representative images of intact human DRG -SC explant (top), immune-stained with peripherin (pink) and Dapi (blue) (mid), or NeuN (yellow) and Dapi (blue) (bottom). (**G)** Representative image of 24 well plate preparation where the DRG is submerged in one well and the nerve root is draped over the adjacent well, and the spinal cord is submerged in the third well ensuring the tissues are separated and no fluid overflows. The fluid was collected from either the DRG well or the spinal cord well every 30 minutes for a total time course of 90minutes. The (1µM) capsaicin was added to the DRG well at the second time-point (30min mark). (**H)** Time course of CGRP release across the DRG (left) and the spinal cord (right) when the DRG was stimulated with 1µM, demonstrating that in the intact DRG-SC preparation, when the DRG was stimulated with 1µM capsaicin, detecting a significant increase in CGRP release in the spinal cord. Data are means ± SEM; n = 4 individual donors; **P < 0.01 versus baseline value at 30-minute timepoint.

For the intact DRG-spinal cord preparation, the DRG remained connected to the spinal cord via intact dorsal roots, with ventral roots transected. Representative imaging confirmed preservation of tissue architecture and afferent connectivity (Figure 1F), and the compartmentalized 24-well configuration ensured physical and biochemical separation of DRG and spinal cord compartments (Figure 1G). Interestingly, stimulation of the DRG with 1 µM capsaicin elicited a robust 1.8-fold increase (180%) in CGRP release exclusively within the spinal cord compartment relative to baseline (100%), with no detectable increase in the DRG well (Figure 1H). These findings demonstrate spatially restricted CGRP release from central terminals of human nociceptors following peripheral TRPV1 activation.

### Nav1.8 activity in the DRG and dorsal root is necessary for central CGRP release

To define the locus of Nav1.8 action on nociceptive transmission in response to TRPV1 stimulation, we applied the selective Nav1.8 inhibitor suzetrigine or broad sodium channel blocker bupivacaine to the DRG in the intact DRG-spinal cord circuit (Figure 2A). Pre-treatment of the DRG cell body with 100µM bupivacaine (25, 26) abolished capsaicin-evoked CGRP release in the spinal cord (Figure 2B). Similarly, selective NaV1.8 inhibition in the DRG cell bodies with 20nM suzetrigine (27, 28) fully blocked spinal cord CGRP release (Figure 2C).

**Figure 2.**
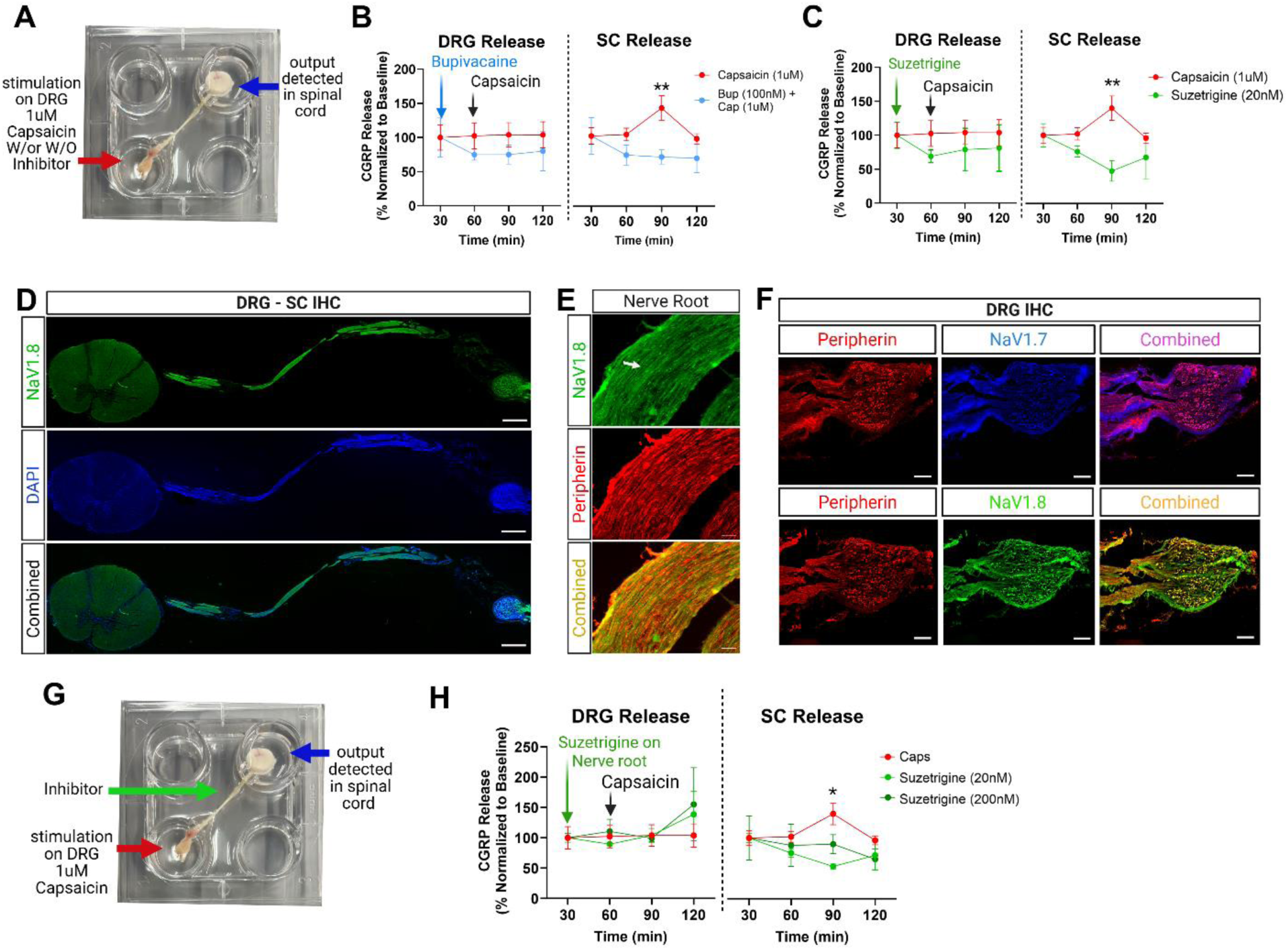
Nav1.8-dependent signal propagation in the DRG and nerve root is necessary for central CGRP release. Selective inhibition of Nav1.8 in the DRG or nerve root is sufficient to abolish central CGRP release, identifying peripheral sensory axons as the critical site of Nav1.8-dependent nociceptive signal propagation. (**A**) Representative image of four well plate preparation where the DRG is submerged in one well and the nerve root is extended across plate and the spinal cord is submerged in the well across ensuring the tissues are separated and no fluid overflows, but the dorsal roots are also submerged. The fluid was collected from either the DRG well or the spinal cord well every 30 minutes for a total time course of 120minutes. The sodium channel inhibitor bupivacaine (100 µM) or selective Nav1.8 inhibitor suzetrigine (20nM) or vehicle were added at the 30-minute timepoint as pre-treatment and then added in conjunction with capsaicin (1µM) at the 60minute mark to the DRG well. (**B)** Time course representing the CGRP release across the DRG (left) and the spinal cord (right), demonstrating that in the intact DRG-SC preparation, when DRG is co-treated with bupivacaine and capsaicin, release of CGRP in the spinal cord is attentuated. (**C**) Time course representing CGRP release across the DRG (left) and the spinal cord (right), demonstrating that in the intact DRG-SC preparation, when DRG is co-treated with suzetrigine and capsaicin release of CGRP in the spinal cord is blocked. Data are means ± SEM; n = 3-4 individual donors. ** P < 0.01 by repeated measures (RM) two-way ANOVA and Tukey’s multiple comparison test **(D**) representative immunohistochemical image of nerve root looking at Nav1.8 (green), DAPI (blue), and merge between the two. Scale bar = 5mm (**E)** Representative 40X magnification image of the nerve root looking at immunoreactivity of Nav1.8 (green), peripherin (red), and the merge (yellow). Scale bar = 100μm (**F)** Representative immunohistochemical images of peripherin (red), Nav1.7 (blue) and the merge of the two (top); peripherin (red), Nav1.8 (green) and the merge of the two (bottom) on hDRG and a portion of the nerve root. Scale bar = 1mm**(G)** Representative image of four well plate preparation where the DRG is submerged in one well and the nerve root is extended across plate and the spinal cord is submerged in the well across ensuring the tissues are separated and no fluid overflows, and the nerve roots are also submerged. The fluid was collected from either the DRG well or the spinal cord well every 30 minutes for a total time course of 120 minutes. Suzetrigine (20nM & 200nM) was added at the 30-minute timepoint as pre-treatment and then added again to the nerve root at the 60-minute mark. Capsaicin (1μM) was added to the DRG well at the 60-minute mark. (**H)** Time course representing the CGRP release across the DRG (left) and the spinal cord (right), demonstrating that in the intact DRG-SC preparation, when nerve root is treated with suzetrigine and the DRG is treated with capsaicin, CGRP release in the spinal cord is blocked. * P < 0.05 by repeated measures (RM) two-way ANOVA and Tukey’s multiple comparison test.

Immunohistochemical analysis revealed robust expression of Nav1.8 along the nerve roots and in DRG neurons, co-localizing with the sensory marker peripherin (Figure 2D-F). Building on these findings, selective application of suzetrigine to the dorsal root axons alone (20 & 200nM) was sufficient to prevent central CGRP release following DRG stimulation (Figure 2G–H). These results identify peripheral sensory axons, including the dorsal root, as a site of Nav1.8-dependent action potential propagation and demonstrate that selective blockade of Nav1.8 at this site effectively prevents central neuropeptide release.

Pre-treatment of the DRG with either bupivacaine or suzetrigine reduced CGRP levels below baseline in both the DRG and spinal cord compartments. In contrast, when suzetrigine was applied selectively to the nerve root, CGRP levels fell below baseline only in the spinal cord compartment, while remaining unchanged in the DRG (Figure 2B-C, H). These findings likely indicate that this ex-vivo preparation exhibits tonic Nav-dependent activity that contributes to basal CGRP release in the dorsal horn.

### Nav1.8 does not regulate presynaptic CGRP release in the spinal cord

Next, we examined whether Nav1.8 directly regulates presynaptic CGRP release in response to capsaicin stimulation within the spinal cord. Spinal cord slices were treated with clinically used analgesics, morphine, bupivacaine, or suzetrigine before and during capsaicin stimulation. Both morphine and bupivacaine significantly suppressed CGRP release, consistent with presynaptic inhibition or blockade of local ion channels (Figure 3A–B). As expected, bupivacaine nearly blocks all CGRP release (∼70%), while morphine only partially but significantly reduces its release (∼45%) as seen by the area under the curve values (AUC). Control experiments with morphine or bupivacaine alone showed no changes in CGRP release across any of the time points. In contrast, suzetrigine had no effect on capsaicin-evoked CGRP release at either 20nM or 200nM concentrations (Figure 3C–D). These findings indicate that Nav1.8 does not contribute to transmitter release at central terminals, distinguishing its role from drugs that act locally within the spinal cord.

**Figure 3.**
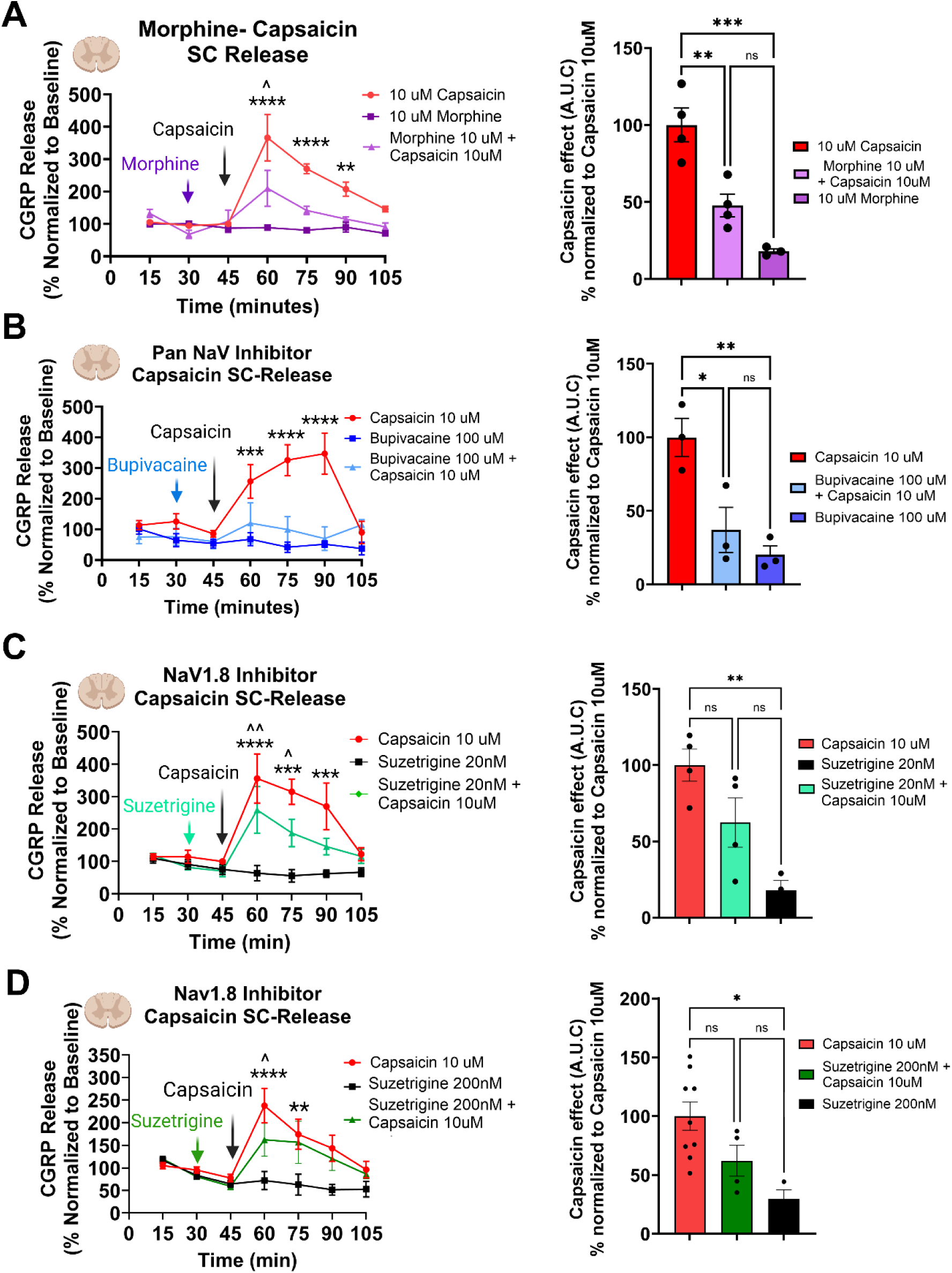
Nav1.8 does not regulate CGRP release through direct action in the human spinal cord. While spinal sodium channel blockade or opioid signaling suppresses CGRP release, selective Nav1.8 inhibition has no effect, demonstrating that Nav1.8 governs peripheral signal propagation rather than central presynaptic release. (**A**) Time course and respective area under the curve (AUC) of CGRP release with 10µM capsaicin and 10µM morphine with and without capsaicin (**B**) Time course of 100µM bupivacaine with and without capsaicin and respective AUC. (**C** and **D**) Time course and respective area under the curve (AUC) of CGRP release with 10µM capsaicin and suzetrigine (20nM & 200nM) with and without capsaicin, demonstrating CGRP evoked release is not prevented by Nav1.8 inhibition in the spinal cord. All AUC are comparison of capsaicin, and respective drugs (morphine, bupivacaine, suzetrigine) with and without capsaicin post 45 min timepoint; data normalized to capsaicin (100%) SEM; n = 3-4 individual donors. *P<0.05, **P<0.01, ***P<0.001 **** P < 0.0001, ^P<0.05, ^^P<0.01, versus same time point in the Vehicle group by repeated measures (RM) two-way ANOVA with Tukey’s multiple comparison test, and AUC by ordinary one-way ANOVA with Tukey’s multiple comparison test.

### Immunohistochemistry confirms Nav1.8-independent spinal release

To validate findings described above with an independent method, we performed quantitative immunohistochemistry on spinal cord slices pre-treated with morphine, bupivacaine, or suzetrigine for 15 minutes and co-treated with 10 µM capsaicin for another 15 minutes before tissue was fresh frozen. Our hypothesis in these experiments was that CGRP release in tissues would appear as diffuse staining throughout the tissue owing to CGRP release from primary afferents whereas non-stimulated tissues would retain prototypical staining for CGRP in the outer lamina of the spinal cord. In vehicle versus capsaicin treated experiments we observed a staining pattern consistent with our hypothesis (Figure 4A); therefore, we examined how engagement of opioid receptors or Nav inhibition would affect CGRP staining in this assay. Morphine and bupivacaine significantly reduced dorsal horn CGRP signal, while suzetrigine produced no detectable effect compared to the capsaicin treated controls (Figure 4B–C). These findings are consistent with our ELISA data demonstrating that Nav1.8 is not directly involved in CGRP release at presynaptic sites of afferent nociceptors in the human dorsal horn.

**Figure 4.**
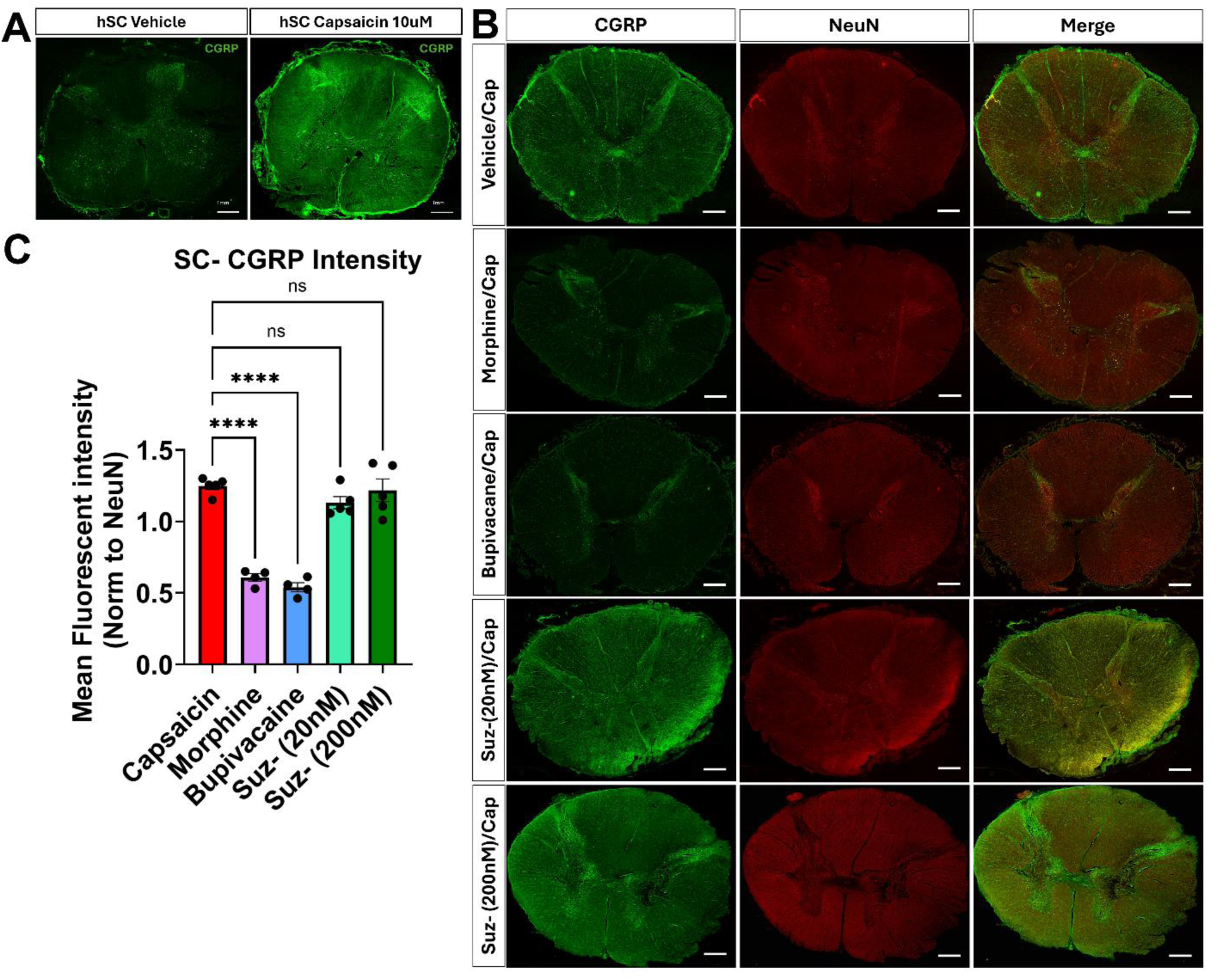
Nav1.8 inhibition fails to suppress CGRP release through a spinal site of action as measured by quantitative immunohistochemistry. Quantitative immunohistochemistry confirms that while morphine and general sodium channel blockade suppress spinal CGRP release, selective Nav1.8 inhibition has no effect. **(A)** Representative immunohistochemical images of CGRP (green) of spinal cord explants treated with vehicle or 10 µM capsaicin. (**B**) Representative immunohistochemical images of CGRP (green) and Neuronal marker (NeuN) on spinal cord explants pre-treated with 10µM morphine, 100µM bupivacaine, 20nM or 200nM suzetrigine or vehicle for 10 minutes, followed by co-treatment with 10µM capsaicin for another 10 minutes, after which all tissues were frozen. Scale bar = 1mm. (**C)** Data for the mean fluorescent intensity of CGRP (green) normalized to NeuN (red) demonstrating morphine and bupivacaine inhibit CGRP release and neither concentration of suzetrigine significantly inhibits CGRP release. Data are means ± SEM; n = 2-3 individual donors with two biological replicates. **** P < 0.0001 versus indicated group by one-way ANOVA with Dunnett’s post hoc test.

## Discussion

In this study, we established a peripheral to central human nociceptive circuit assay to define how peripheral nociceptor activation is converted into central neuropeptide release and to identify the anatomical site of action of the Nav1.8 inhibitor suzetrigine. Our findings address a central mechanistic question raised by the clinical development of Nav1.8 inhibitors: whether selective Nav1.8 blockade is sufficient to suppress central nociceptive output, and if so, at what anatomical locus. Our key findings are: 1) that Nav1.8 inhibition along nociceptor axons is sufficient to block CGRP release in the spinal cord when those axons are activated by capsaicin; and 2) that Nav1.8 blockade in the spinal cord is insufficient to attenuate capsaicin-evoked CGRP release. These findings suggest that Nav1.8 is critical for carrying action potentials when human nociceptors are activated, consistent with previous patch clamp electrophysiology studies (12, 14–16).

Measurement of evoked SP and CGRP release has been utilized for decades in animal models to quantify primary afferent excitability and spinal neurotransmission (29–32). We therefore selected CGRP as the functional readout, given its abundant expression in TRPV1-positive nociceptors and the ability to reliably evoke its release using the well-validated TRPV1 agonist capsaicin (33, 34). Using human spinal cord slices in addition to DRG-SC tissues enabled us to distinguish outcomes between interruption of axonal action potential propagation and modulation of presynaptic neurotransmitter release at central terminals. Morphine suppressed capsaicin mediated CGRP release in spinal cord but still allowed incremental release above baseline, consistent with its presynaptic inhibition of calcium-dependent exocytosis, whereas bupivacaine nearly abolished release, reflecting blockade of terminal sodium channels (Figure 3A-B) (35, 36). These pharmacologic effects closely mirror clinical observations where morphine provides analgesia without sensory blockade, and local anesthetics produce profound suppression of nociceptive signal propagation (37–41). The consistency between the clinical outcomes and the physiological response in our experiments supports the translational relevance of the human spinal cord explant model.

Strikingly, the selective Nav1.8 inhibitor suzetrigine had no effect on spinal CGRP release (Figure 3C-D) when applied directly on the spinal cord slices. The results are consistent with the immunohistochemistry findings showing Nav1.8 protein is absent from the spinal cord but is robustly expressed in DRG neurons and dorsal roots (Fig 2D-F). In contrast, other sodium channels such as Nav1.7 is expressed within superficial dorsal horn laminae, providing a mechanistic explanation for the differing effects of nonselective sodium channel blockade with bupivacaine (42). With this in mind, we decided to test suzetrigine in our intact human DRG-spinal cord circuit preparation. Capsaicin-induced TRPV1 activation, at the DRG level, triggers CGRP release within the spinal cord, which was blocked by pre- and co-treatment of bupivacaine with capsaicin on the DRG. This demonstrates that the model preserves classic peripheral-to-central signal propagation, *a first* with ex-vivo human tissue to be presented. Selective inhibition of Nav1.8 in the DRG or dorsal root abolished capsaicin-evoked CGRP release centrally, demonstrating that peripheral Nav1.8 activity is sufficient to couple nociceptor activation to central neuropeptide (Fig 3). These findings suggest Nav1.8 plays a pivotal role in propagation of action potential from DRG to the spinal cord.

Our findings have implications for development of next generation Nav1.8 inhibitors. Clinical trials demonstrate that suzetrigine is efficacious for acute pain when administered systemically (13, 17, 18), yet no studies have evaluated intrathecal Nav1.8 inhibition in humans. Prior to this work it remained unclear whether Nav1.8 localizes along centrally projecting human DRG axons or contributes directly to neurotransmitter release at central terminals. Our findings clarify this anatomical uncertainty, showing that Nav1.8 supports action potential propagation along DRG axons and dorsal roots but does not regulate neurotransmitter release locally within the dorsal horn. These results suggest that increasing CNS penetration of Nav1.8 inhibitors is unlikely to enhance efficacy. Notably, while multiple reviews mention a low brain to plasma ratio for suzetrigine, our review of the literature and publicly available FDA documents did not find any direct measurement of CNS penetration for suzetrigine in humans (27, 43, 44)

Some studies in the past have proposed the concept of intraganglionic signaling in regard to sensory ganglia (45, 46). In our studies with intact circuit, post capsaicin stimulation of DRG neurons, we did not detect CGRP from the supernatant collected from DRG, while CGRP release was clearly observed in the spinal dorsal horn. This finding suggests that in intact adult human circuits, neuropeptide output is tightly compartmentalized to central terminals. This contrasts with acutely dissociated DRG tissue or axotomized DRG cultures, where somatic CGRP release is readily observed, and underscores the importance of preserving axonal continuity when interrogating human nociceptive biology (47). Although intraganglionic CGRP signaling has been proposed, whether human DRG neurons release neuropeptides somatically under physiological conditions remains unresolved. Additional research is needed to ascertain whether paracrine CGRP, or other neuropeptide, signaling takes place within the sensory ganglia; however, our data suggest the extent of release is minimal to be detectable in the supernant post capsaicin stimulation.

An interesting question arises regarding the observed clinical effects of suzetrigine and the extent of inhibition of action potential propagation in our preparation. Although local application of suzetrigine to the DRG and spinal nerve root nearly abolished capsaicin-evoked CGRP release in the spinal cord, more modest clinical efficacy has been reported with oral administration in acute and chronic pain settings (48, 49). This discrepancy could reflect insufficient concentration achieved at the level of the DRG cell bodies or proximal nerve root following systemic administration to fully inhibit Nav1.8-dependent action potential propagation. This limitation could be addressed clinically via perineural application of suzetrigine or the development of a Nav1.8 inhibitor with increased nerve bundle penetration.

More broadly, this work establishes that peripheral to central human nociceptive circuits can reveal mechanistic features of pain transmission that are obscured in dissociated neurons and in isolated spinal cord preparations. By preserving native anatomy and directional connectivity, these human explants allow direct localization of drug action within the nociceptive axis and provide a translationally relevant system for evaluating human-specific analgesic mechanisms.

Several limitations merit consideration. Our model reflects acute TRPV1 driven activation rather than chronic pain states and lacks systemic immune and glial influences present in vivo. CGRP release represents a central neuropeptide output but does not fully capture nociceptive transmission, which also depends on glutamate release and postsynaptic activation. The acutely isolated explant preparation may alter neuronal properties, and our sample size was not powered to detect sex or age specific differences.

## Methods

### Sex as a biological variable

Our study examined male and female human tissues, and similar findings are reported for both sexes.

### Human DRG and spinal cord recovery

All human tissue procurement procedures were approved by the Institutional Review Board at the University of Texas at Dallas. Human DRGs and spinal cord were surgically extracted using a ventral approach (50) from organ donors within 3 hours of cross-clamp and placed immediately on artificial cerebrospinal fluid (aCSF). All tissues were recovered in the Dallas area via a collaboration with the Southwest Transplant Alliance as described in a previously published protocol Tissue samples from 21 donors were used in this study, information is provided in Table 1.

**Table 1.** Donor Demographic.

| Donor ID | Sex | Age | COD | Race/Ethnicity |
| --- | --- | --- | --- | --- |
| UTD-DN0344 | M | 50 | Anoxia | White |
| UTD-DN0346 | F | 42 | Drug OD | White/Hispanic |
| UTD-DN0338 | F | 41 | CVA/Stroke | White |
| UTD-DN0339 | M | 30 | Head trauma/MVA | White |
| UTD-DN0336 | F | 41 | Anoxia | white |
| UTD-DN0340 | F | 43 | Anoxia | White/Hispanic |
| UTD-DN0341 | M | 59 | GSW | white |
| UTD-DN0348 | M | 25 | MI | Black |
| UTD-DN0334 | M | 20 | Anoxia/Cardiovascular | White |
| UTD-DN0270 | F | 62 | Anoxia | white |
| UTD-DN0379 | F | 44 | CVA/Stroke | White |
| UTD-DN0380 | M | 32 | Head Trauma/MVA | White |
| UTD-DN0386 | M | 19 | GSW | White/Hispanic |
| UTD-DN0392 | F | 34 | Respiratory | white |
| UTD-DN0399 | F | 24 | OD | white |
| UTD-DN0411 | F | 25 | Anoxia | Black |
| UTD-DN0412 | M | 71 | Anoxia | White |
| UTD-DN0381 | F | 45 | Heart Attack | White |
| UTD-DN0416 | M | 48 | stroke/ CVA | white |
| UTD-DN0422 | F | 27 | GSW | white |
| UTD-DN0459 | M | 64 | Stroke/ CVA | White |

\*COD: Cause of Death, GSW: Gunshot wound, MVA: Motor vehicle accident, OD: Overdose, Anoxia: Oxygen deprivation, Stroke: Cerebrovascular accident.

### Spinal cord and DRG dissection

Spinal cord meninges were removed, and the pia was cleaned as much as possible without disrupting the integrity of the tissue in aCSF on ice. We chose to use a set of sharp scissors (F.S.T, 14060-10) to make a clean fast cut to prevent any destruction of the tissue and used a ruler to measure 2mm of tissue before making the cuts. For the DRG, the tissue was cleaned down to the bulb and then placed inside a 1.0mm mouse brain block and cut using razor blades. Each cut section was immediately placed into Tyrode’s buffer (NaCl 119 mM, KCl 2.5 mM, MgCl2 2 mM, CaCl2 2 mM, HEPES 25 mM, Glucose 30 mM) and placed in a shaking incubator for the initial equilibration time. DRG tissue sections were placed in an 8-well chamber slide (400ul), and the spinal cord sections were placed in 24-well plates (600ul). Fluid was removed and replenished every 30 minutes for up to two hours before the experiment started. For the intact DRG and spinal cord preparation, the meninges were carefully removed, and each individual root was slowly separated. The ventral root connection was severed and only the dorsal root connections were kept. The spinal cord is sectioned into 2mm cuts with at least 3 dorsal roots left intact. The DRG was lightly cleaned but kept intact for the most part. Tissues that were used for IHC were also equilibrated for two hours and then incubated with the respective inhibitor and/or agonist for 15 minutes and immediately frozen in pulverized dry ice.

### Immunohistochemistry

The human spinal cord explants were gradually embedded in OCT in a cryomold by adding small volumes of OCT over dry ice to avoid thawing. All tissues were cryostat sectioned at 20 μm onto SuperFrost Plus charged slides. Slides were removed from the cryostat and immediately transferred to 4% PFA for 15 minutes. The tissues were then dehydrated in 50% ethanol (5 min), 70% ethanol (5 min), 100% ethanol (5 min), 100% ethanol (5 min) at room temperature. The slides were air dried briefly and then boundaries were drawn around each section using a hydrophobic pen (ImmEdge PAP pen, Vector Labs). When hydrophobic boundaries dried, the slides were submerged in blocking buffer (10% Normal Goat Serum, 0.3% TritonX 100 in 1X Phosphate Buffer Saline (PBS)) for 1 hour at room temperature. Slides were then rinsed in 1x PBS, placed in a light-protected humidity-controlled tray and incubated in primary antibodies anti-CGRP (1:300 Immunostar, 24112), Anti-NeuN (1:500 MilliporeSigma, MAB377), Anti-Peripherin (chicken 1:1000; EnCor, AB_2284443), anti-Nav1.8 (1:300 NeuroMab N134/12) diluted in blocking buffer overnight at 4°C. The next day, slides were washed in 1X PBS and then incubated in their respective secondary antibody Alexa Fluor goat anti-mouse IgG 555 (1:2000; Invitrogen A-21449, lot 2186435), Alexa Fluor goat anti-rabbit IgG 488 (1:2000; Invitrogen A-11034, lot 2110499), Alexa Fluor goat anti-chicken IgG 647 (1:2000; Invitrogen A-21449, lot 2186435), with DAPI (1:5000; Cayman Chemical; Cat # 14285) diluted in blocking buffer for 1 hour at room temperature. Sections were then washed in 1X PBS, air dried and cover slipped with Prolong Gold Antifade reagent. Whole slides were imaged at 20X magnification using an Olympus VS200 research slide scanner. For quantification, regions of interest (ROIs) were defined encompassing the superficial laminae (I/II) of the dorsal horn. The mean fluorescent intensity (MFI) of CGRP was measured bilaterally and normalized to the MFI of NeuN within the same ROI to account for variations in neuronal density and sectioning.

Background corrected values were averaged per tissue section for statistical analysis

### CGRP release

For the DRG and spinal cord explants alone, after the 2-hour equilibration time, the fluid is switched to fresh Tyrode’s buffer with added thiorphan DL (16uM, Santa Cruz CAS 76721-89-6). Media was collected every 15minutes for a total of 90 minutes (Capsaicin only experiments) or 105 minutes (inhibitor with capsaicin experiments). The capsaicin was added at the 30-minute mark for the dose response experiments both in the DRG and the spinal cord. For experiments where there was an inhibitor on board (Morphine, Hikma 0641-6132-25) (Suzetrigine, MedChemExpress HY-148800) or (Bupivacaine, Sigma-Aldrich 73360-54-0), the respective inhibitor was added at the 30-minute mark, and then capsaicin with the inhibitor was added at the 45-minute time point. The fluid collected was then analyzed for amounts of CGRP released using manufacturer recommendations for a human CGRP EIA kit (Cayman Chemicals 589101). The data is shown as a percentage change, and each individual tissue was normalized to its own 2-point baseline release.

### Viability Assays

CellTiter 96 Aqueous one solution cell proliferation assay (MTS). Each spinal cord or DRG section was preincubated with 20ul of the dye per 100ul of Tyrode’s buffer for 2 h at 37°C in the shaking incubator. The fluid is then removed and transferred to a 96-well plate divided into triplicates and then the absorbance at 490 nm was recorded. This served as the Pre-treatment value. Various concentrations of capsaicin or vehicle were used to treat the spinal cord (1µM, 5µM, 10µM) and DRG (0.5µM, 1µM), for a total of 30 minutes. The fluid is then removed, and the tissue is once again incubated with the MTS (20µl of the dye per 100µl of Tyrode’s buffer) for 2 h at 37°C and this serves as the post-treatment MTS value. Fluid was transferred to a 96-well plate divided into triplicates and then the absorbance at 490 nm was recorded. The percent change in viability was determined using the previously published formula: ((post-treatment MTS – Pre-treatment MTS)/Average Pre-treatment MTS)) *100

### Statistics

Data was presented as the mean ± SEM. All analysis was done using GraphPad Prism 10.6.1. We used a One-way ANOVA and Tukey’s multiple comparison test for the viability assays and the AUC. For the IHC analysis, we used a One-way ANOVA with Dunnett’s post hoc test. For all of the time courses, we used a two-way ANOVA with Tukey’s post hoc multiple comparison test. P < 0.05 was considered statistically significant.

## Study Approval

Human tissue procurement procedures were approved by the Institutional Review Board at the University of Texas at Dallas (MR-15-237). The Southwest Transplant Alliance (STA) obtained informed consent for research tissue donation from first-person consent or from the donor’s legal next of kin. Policies for donor screening and consent are those established by the United Network for Organ Sharing (UNOS). STA follows the standards and procedures established by the US Centers for Disease Control (CDC) and are inspected biannually by the Department of Health and Human Services (DHHS). The distribution of donor medical information is in compliance with HIPAA regulations to protect donor privacy.

## Data Availability

All data is available in the supporting data.

## Author contributions

A.P, T.J.P, S.M.P, and K.A.G all conceived the study, designed the experiments and co-wrote the manuscript. S.M.P and K.A.G performed the experiments, analyzed the data, and contributed equally to this work. The order of co–first authors was determined by mutual agreement, reflecting the primary site of experimental work and overall project leadership. A.C, P.H, T.K, S.S and GF organized and performed tissue retrieval from human organ donors. I.K, S.R performed the immunohistochemistry experiments and A.K performed the ELISA’s. All authors saw and approved the manuscript. T.J.P and A.P received funding to support the experimental work performed.

## Acknowledgment

The authors thank the organ donors and their families for their enduring gift, as well as our partners at Southwest Transplant Alliance without whom this research would not be possible. The authors would also like to thank the donor recovery teams in the Price lab at UT Dallas.

